# Using Large Language Models to Assemble, Audit, and Prioritize the Therapeutic Landscape

**DOI:** 10.1101/2025.11.14.688562

**Authors:** Anugraha Thyagatur, Nithin Sonti, Rahul Vijayan, Ayaan Parikh, Roland Faller

## Abstract

We present an AI-assisted pipeline for disease-specific drug landscape analysis. Given a disease name, the system assembles a comprehensive, evidence-based view of therapeutic assets by integrating structured sources (such as ClinicalTrials.gov and ChEMBL) and unstructured sources (such as publications, press releases, and patents). Large language models are used in a constrained, auditable mode to normalize drug aliases, resolve drug–target/mechanism of action annotations, and harmonize program status across records. The output is a disease-centric map that spans preclinical assets, not-yet-approved assets (both active and discontinued/shelved), and FDA-approved drugs suitable for re-purposing. Assets are ranked using interpretable, evidence-based scoring heuristics that combine trial volume and clinical phase, endpoint outcomes, biomarker support, recency of activity, and regulatory designations, along with penalties for safety signals and non-pharmaceutical interventions, as well as proportional adjustments for operational versus scientific discontinuations. Case studies in Alzheimer’s disease, pancreatic cancer, and cystic fibrosis demonstrate generality, coverage, and discrimination across mechanisms and stages. This framework provides a transparent method to assemble and prioritize the therapeutic landscape for any disease, unifying disparate data into a coherent and analyzable representation.

## 2 Introduction

A comprehensive and contemporary overview of the emerging therapeutic landscape is essential for accelerating clinical translation. It provides the strategic intelligence needed to pivot R&D pipelines toward the greatest unmet needs, benchmark novel assets against competitors, and optimize trial design, ultimately reducing the time and cost of bringing effective therapeutics to patients. Such an overview must enumerate active investigational/shelved drugs across all clinical phases, identify programs paused for non-scientific reasons, highlight preclinical candidates with validated mechanisms of action, and capture drug re-purposing opportunities involving therapies approved for other indications [1, 2, 3, 4, 5]. In practice, constructing this view is a significant informatics challenge, as the requisite evidence is fragmented across disparate sources, such as clinical trial registries, regulatory approval databases, and the scientific literature [6].

Each repository, from clinical trial registries to the scientific literature, provides only a partial account of a therapeutic asset’s history, which complicates any attempt to form a complete and accurate understanding. This fragmentation creates significant barriers to data integration [7, 8]. For instance, a single drug is often identified by different names or development codes across various databases, making the simple task of tracking one asset’s complete journey a complex data-linking problem [9, 10, 11, 12]. Compounding this issue, critical details such as the current status of a trial or the specific reasons for its discontinuation are frequently recorded in inconsistent, free-text narratives instead of standardized fields, making any systematic or automated analysis of trial outcomes extremely difficult [13, 14, 15, 16].

These disconnected information sources, combined with the high attrition rate in drug development, underscore the need for an integrated framework. Drug development is arduous: nearly 90% of candidates entering clinical testing do not ultimately achieve U.S. Food and Drug Administration (FDA) approval, contributing to a large and ever-changing pool of investigational assets [17, 18]. Programs are discontinued for diverse reasons, ranging from insufficient efficacy to portfolio reprioritization, and such rationales are often reported unevenly [13]. In this environment, an expert attempting to assemble a reliable landscape must reconcile disparate identifiers, normalize inconsistent status narratives, and integrate data from preclinical, clinical, and cross-indication studies—tasks that are labor-intensive when performed manually.

Several commercial platforms attempt to provide consolidated views of the drug development pipeline [19, 20, 21]. While valuable for certain business applications, many proprietary platforms require substantial licensing fees and are not open-source. These cost and access constraints limit adoption by academic labs, non-profits, and smaller organizations, and reduce opportunities for community-driven extension and reuse. Open-source academic efforts have primarily focused on data consolidation and entity resolution. For example, CDEK [22] is a valuable public knowledge base that curates and disambiguates entities (drugs, organizations) from structured sources such as ClinicalTrials.gov and other drug databases. However, its main objective is to create a comprehensive catalog for lookup. A significant benefit can be provided by a tool that moves beyond aggregation to perform evidence-weighted prioritization of the drug development pipeline, thereby identifying the most promising therapeutic assets. This demands an adaptable framework that can parse, analyze, and score assets based on the scientific context found in publications, press releases, and clinical reports.

Large Language Models (LLMs) offer a practical means to overcome these specific informatics barriers. They can directly address the “complex data-linking problem” by analyzing text to link disparate aliases and development codes to canonical entities. Furthermore, they can normalize “inconsistent, free-text narratives” (such as those describing discontinuation reasons or trial outcomes) into standardized, analyzable fields, thereby automating the extraction of salient facts from fragmented sources. Crucially, when employed as reasoning agents with access to tools like web search, LLMs are not limited to their static training data. They can actively query for and retrieve the most recent, real-time information from publications, press releases, and trial updates. This allows the system to analyze and reason over the latest evidence, ensuring the resulting landscape is both comprehensive and contemporary.

In this work, we introduce an open-source informatics system that addresses the above challenges by constructing a unified, auditable landscape of emerging therapeutics for any specified disease. The system integrates heterogeneous data sources and provides a transparent, end-to-end, disease-specific view of investigational drugs and their supporting evidence. Given a disease query, it aggregates and deduplicates asset mentions across clinical trial registries, regulatory lists, curated compound and target databases, and the literature (Europe PMC [23], PubMed [24]). It then harmonizes identifiers and program metadata and normalizes discontinuation rationales. Next, it assembles a disease-centric map that spans preclinical leads, active investigational agents, paused programs, and cross-indication drugs under investigation. Finally, it prioritizes not-yet-approved assets using an interpretable, evidence-weighted heuristic while retaining full source provenance for expert review. To demonstrate the generality of this framework, we apply it to case studies in Alzheimer’s disease, pancreatic cancer, and cystic fibrosis, diseases spanning neurodegenerative, oncologic, and genetic domains, respectively. These examples showcase the system’s ability to adapt its integrative approach across distinct therapeutic areas, underscoring its broad applicability as an open-source tool for the scientific community.

## 3 Methods

### 3.1 System Architecture and Workflow

We implemented a modular informatics pipeline that constructs a comprehensive therapeutic landscape for a specified disease, as shown in **Figure 1**. The system operates through a central orchestrating agent that executes a multi-stage workflow. This workflow begins with systematic evidence aggregation from disparate sources and proceeds through stages of data harmonization, multi-modal enrichment, and evidence-based scoring to identify and prioritize promising therapeutic assets. All external data requests are executed via a rate-limited HTTP client with a persistent caching layer to ensure reproducibility and optimize API resource management.

**Figure 1:**
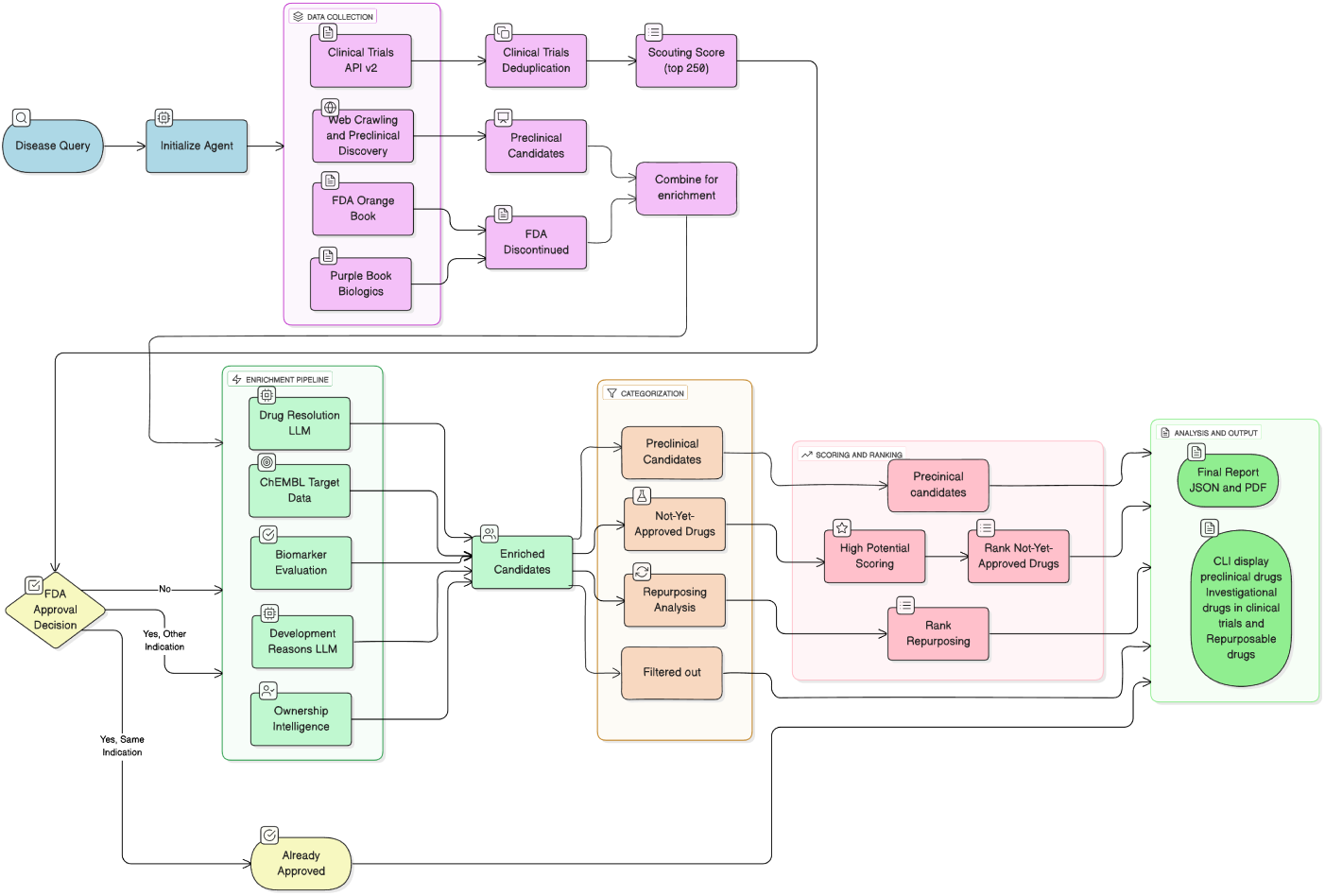
AI-assisted triage pipeline. Data collection, enrichment, and categorization gathers disease-scoped evidence, normalizes entities, assigns status/reasons, audits endpoints/biomarkers/safety, and attaches provenance. Scoring and ranking partitions assets and computes the final score S for investigational programs, with a repurposing branch for FDA-approved drugs not indicated. Analysis and output compiles evidence dossiers and ranked lists for expert review.

### 3.2 Stage 1: Data Collection and Initial Scoring

The first stage of the pipeline focuses on collecting and consolidating all potential therapeutic candidates mentioned in connection with the target disease.

A dedicated web-crawler discovers preclinical candidates not yet in clinical trials through a multi-stage validation pipeline. The system begins by performing targeted web searches using LLM-powered queries such as ‘disease novel compound preclinical’ with exclusion terms (-clinical, -phase, -trial) to filter out clinical trial content, and ‘disease investigational new drug IND preclinical’ to identify research papers, patents, and company announcements containing specific compound names and research codes in the preclinical development stage. The LLM extracts compound identifiers from retrieved web content, focusing on specific naming patterns such as research codes (company-prefix-number format like RO-1234, BMS-456), chemical names (actual drug names like temozolomide, imatinib), and antibody designations (anti-target formats) rather than generic drug class descriptions like ‘KRAS inhibitor’ or ‘monoclonal antibody. Each extracted compound undergoes a two-tier validation process. First, database validation verifies compound existence through parallel searches of the PubChem and ChEMBL APIs, with compounds receiving validation scores (0.5 points for a PubChem match and 0.5 points for a ChEMBL match), requiring a minimum threshold of 0.5 to pass this validation step. Second, the clinical trials filtering step queries the ClinicalTrials.gov database to ensure the system identifies only genuinely early-stage opportunities, confirming that no existing trials for the compound are registered for the target disease. Only compounds that pass all validation stages and show no clinical trial activity for the target disease are classified as preclinical candidates, ensuring that the system identifies genuinely early-stage therapeutic opportunities.

, Additionally, the FDA Orange Book and Purple Book databases are queried to identify approved, discontinued, and withdrawn drugs. However, the databases do not include safety-withdrawn determinations or FDA recall data. This process identifies a critical pool of assets that may be valuable for drug repurposing, even if they are not currently marketed.

**, Orange Book (, Small Molecules):** The system parses the Orange Book products.txt file, which contains a market status field with three possible values: RX (prescription), OTC (over-the-counter), or DISCN **(discontinued)**. Products marked with DISCN are automatically classified as discontinued drugs.

**, Purple Book (, Biologics):** For biological products, the system queries the Purple Book database, which maintains similar status fields indicating whether biosimilars and reference biologics are currently marketed or have been discontinued.

**, Drugs@, FDA Integration:** The system cross-references Orange/, Purple Book entries with FDA labeling to confirm disease relevance. Each asset is linked to its label using application identifiers (, NDA/, ANDA/, BLA) when available or by normalized ingredient, dosage form, and route when identifiers are absent. Indication evidence is extracted from the Indications and Usage section of the label (via the open, FDA Drug Label API indications usage field), with Drugs@, FDA consulted for any gaps. An asset is retained as a discontinued/not-marketed candidate if the corresponding application is marked as withdrawn or terminated (small molecules) or if the Purple Book license is inactive/withdrawn (biologics) and the label text supports the target disease.

, Next, candidates from clinical trials are accessed through the ClinicalTrials.gov v2 REST API, which is queried using a systematic multi-step process. The system initializes a ClinicalTrialsClient that interfaces with the API base URL HTTP://clinicaltrials.gov/api/v2 and employs rate-limited HTTP requests with caching to manage API quotas.

**Query Construction:** The client constructs API queries using v2-specific parameters, with disease conditions specified via query.cond and interventions via query.intr, while status and phase filters utilize pipe-separated values (filter.overallStatus and filter.phase). For comprehensive disease-based discovery, the primary query searches all trials for a given condition using await self.search trials(condition=disease, max results=20000).

**Pagination and Data Extraction:** The API responses are paginated with a maximum page size of 1000 studies, requiring iterative requests using pageToken parameters until all relevant trials are retrieved. For each study, the client extracts structured data from the protocol section, including NCT ID, trial title, overall status, study phases, interventions, enrollment numbers, start/completion dates, termination reasons (why stopped), and sponsor information from nested modules (identification Module, status Module, design Module, arms Interventions Module, sponsor Collaborators Module).

**Response Processing and Data Consolidation:** The extracted trial records are first aggregated by therapeutic intervention, with each drug’s trial history consolidated to calculate statistics, including total trials, completion rates, failure patterns, and phase progression. Following this initial aggregation, a deduplication and consolidation process is performed on all clinical trials for the given disease. Asset records are grouped using normalized drug names that undergo comprehensive cleaning, including the removal of salt forms (hydrochloride, sodium, sulfate, etc.), dosing information, administration routes, and parenthetical content, followed by case-insensitive and whitespace-normalized string matching. Records sharing a canonical identifier are then merged by summing their trial counts, combining their unique sponsor sets, and preserving the metadata from the record corresponding to the most advanced development phase. For example, separate trial records for ‘Donepezil Hydrochloride 10 mg’, ‘donepezil hcl’, ‘DONEPEZIL (oral tablet)’, and ‘Donepezil Sodium’ would all be normalized to ‘donepezil’ and merged into a single consolidated entry with combined statistics from all naming variants. Each consolidated candidate record is tagged with its source(s) to maintain full provenance, forming the foundation for subsequent scoring and analysis.

#### 3.2.1 Scouting Score Calculation

The aggregated clinical trial data are then used to calculate an initial Scouting Score for each asset. This composite score is built in several layers to form a comprehensive, evidence-based heuristic, as shown in Figure S1 and equation 1.

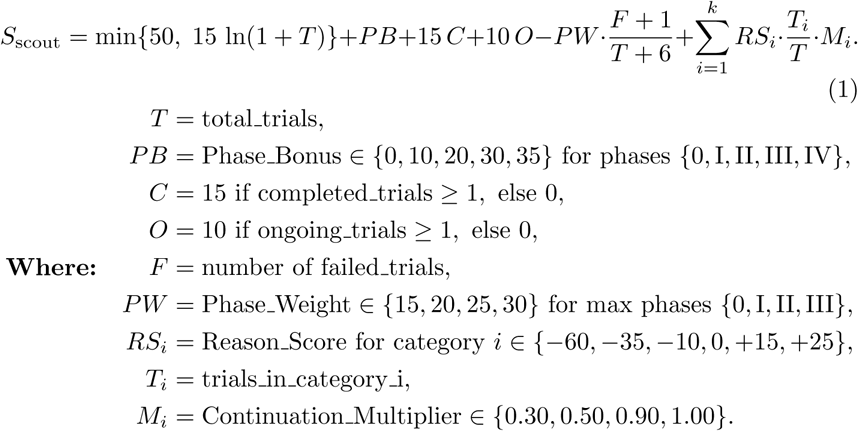

The foundation of the score is the Trial Points metric, which uses a logarithmic function, min(50, 15×ln(1+total trials)), to award points for trial volume with diminishing returns, capped at a maximum of 50. To this base, several positive modifiers are added: a Phase Bonus of up to 35 points for assets reaching late-stage trials (Phase IV), with 30 points for Phase III, 20 points for Phase II, and 10 points for Phase I assets. A 15 point Completion Bonus for assets with at least one completed trial and a 10 point Ongoing Bonus for programs with currently active trials.

Conversely, a Failure Penalty is subtracted, calculated by multiplying a phase-dependent weight by a smoothed failure rate. The phase-dependent weights are 30 for Phase III failures, 25 for Phase II failures, 20 for Phase I failures, and 15 for assets that have not progressed beyond preclinical studies. This smoothed failure rate, (failed trials+1)/(total trials+6), incorporates a Beta(1,5) Bayesian prior that assumes a baseline failure probability. This prevents the over-penalization of drugs with limited trial data that would occur when using raw failure rates, where a single early failure could result in a 100% failure rate and maximum penalty, even though the drug may have significant therapeutic potential but is simply understudied with only one failed trial.

Finally, Reason Adjustments are applied using a two-stage process that distinguishes between scientific failures and operational discontinuities. First, at the individual trial level, the system classifies the official termination reason provided in the ‘why stopped’ field of each terminated trial. The preferred method employs an LLM (gpt-4o) which, in addition to the trial’s ‘why stopped’ text, uses web search to gather external context from press releases, news articles, and regulatory filings. This allows for a robust classification of each trial into one of seven categories, each with a base score (Safety: −60, Efficacy: −35, Regulatory: −10, Protocol: +15, External (COVID-19, natural disasters, war, etc): +15, Business: +25, Recruitment: +25, Others: 0).

Second, at the aggregate drug level, these individual trial classifications are consolidated. The final score adjustment for a drug is calculated proportionally based on the fraction of its total trials that fall into each category. For example, if an asset with 10 total trials had two terminated for safety reasons, the adjustment from that category would be −12 points (20% of the −60 base penalty). This proportional approach ensures that drugs halted primarily for operational reasons receive a net positive adjustment, while those with a consistent pattern of scientific failure are appropriately penalized. Furthermore, the LLM’s web search also identifies whether drugs with safety or efficacy failures are still in active development (e.g., reformulated or pivoted). If so, their penalties are significantly reduced to reflect continued investment in the asset. For the above asset, with 2 terminated trials out of 10 total trials, the LLM’s pipeline analysis can significantly reduce safety and efficacy penalties based on continued development status. If the company has reformulated the drug by modifying the formulation, dosing, or delivery method to address the specific issues that caused previous failures, the penalty is reduced by 70% (from −12 to −3.6 points). If the institution/sponsor is continuing the development of the original drug formulation for the same indication, showing persistence despite setbacks but without addressing the root causes, the penalty is reduced by 50% (from −12 to −6 points). However, if the institution has pivoted to a different indication, essentially abandoning the original disease target to test the drug for a completely different condition, the penalty is reduced by only 10% (from −12 to −10.8 points). These reductions reflect the continued confidence in the underlying therapeutic potential, despite previous setbacks, with the largest reduction granted to drugs for which companies have actively learned from and addressed the root causes of failure, rather than simply persisting with the same approach or abandoning the original therapeutic application.

The output of Stage 1 comprises three distinct candidate pools: clinical assets retrieved from ClinicalTrials.gov (limited to the top 250 compounds ranked by Scouting Score), preclinical candidates identified through systematic web crawling, and discontinued therapeutic agents extracted from the FDA Orange Book and Purple Book databases. The Preclinical candidates and FDA-discontinued assets are consolidated into a unified pool. The top-ranked clinical assets undergo regulatory status validation through FDA Approval Decision triage. Assets with existing approval for the target indication are directly routed to the final analytical output. Clinical assets lacking FDA approval or possessing approval for alternative indications are merged with the consolidated preclinical and discontinued asset pool. This integrated candidate ensemble subsequently advances to the Enrichment Pipeline for comprehensive characterization.

### 3.3 Stage 2: Multi-Modal Enrichment and Characterization

In the second stage, each candidate from stage 1 undergoes a comprehensive enrichment process through five parallel processing modules to build a holistic evidence profile(see (**Figure 1**)).

#### 3.3.1 Drug Resolution LLM

A batch-processing resolver powered by GPT-4o standardizes all asset identifiers. This critical step converts internal research codes and synonyms into canonical drug names through systematic queries to the ChEMBL database API, PubChem REST API. The system assigns ChEMBL identifiers where available, extracts PubChem compound identifiers (CIDs), and annotates primary molecular targets and mechanisms of action through LLM-powered analysis with a real-time web search that accesses pharmaceutical company press releases, industry news articles, regulatory filings, and publicly available scientific literature(Europe PMC). Additionally, this module identifies FDA special designations, including Fast Track, Breakthrough Therapy, Priority Review, and Orphan Drug status, by analyzing current regulatory announcements and company disclosures. Finally, this module identifies **regional regulatory approvals (FDA, EMA, PMDA, NMPA, Health Canada)**.

#### 3.3.2 ChEMBL Target Data

The system queries ChEMBL’s comprehensive bioactivity database to extract validated molecular targets, bioactivity profiles, and mechanism-of-action annotations. This module cross-references compound identifiers to retrieve high-confidence target associations and pharmacological classifications that inform the assessment of therapeutic potential.

#### 3.3.3 Biomarker Evaluation

The biomarker evaluator employs a systematic analysis to assess the therapeutic efficacy markers. **Step 1: Disease Biomarker Identification** - The system uses GPT-4o with web search capabilities to identify the main biomarkers and surrogate endpoints used to judge treatment effects in the target disease, classifying each as target engagement (PK-PD), efficacy, or safety, while determining the beneficial direction (increase/decrease) for each. **Step 2: Published Results Extraction** - The module searches published literature, conference abstracts, regulatory documents, and press releases to extract quantitative biomarker data. Structured prompts request key data for each biomarker: arm(s) compared, baseline value, change or % change, between-group difference, p-value/CI, and timepoint. Each result must be validated with verbatim quotes and source URLs. **Step 3: Biomarker Effect Rating (BER) Scoring** - Each biomarker outcome receives a Biomarker Effect Rating (BER) on a standardized −2 to +2 scale: +2 for statistically significant improvement, +1 for positive trends without significance, 0 for no clear change, −1 for negative trends, and −2 for statistically significant adverse effects or clinically meaningful safety worsening (e.g., ALT/AST ≥3×ULN, QTc prolongation). **Step 4: Combined Score Calculation** - Individual BER scores are converted to a 0-1 scale using the transformation (*BER* + 2)*/*4. The final score is a strategic average of normalized efficacy biomarkers, but critically, severe safety signals (BER = −2) are also included in the average to penalize dangerous drugs.

**Precision Medicine Defaults:** When quantitative biomarker data is unavailable, the system assigns evidence-based default scores: 0.75 for drugs with precision medicine approaches and 0.65 for those with patient selection strategies. Patient selection is determined through keyword analysis, including “biomarker positive,” “selected patients,” “enriched population,” “amyloid positive,” and “tau positive”.

For example, an Alzheimer’s drug: amyloid reduction (30%, p*<*0.01) receives BER +2 → normalized 1.00; cognitive improvement (15%, p=0.08) receives BER +1 → normalized 0.75; liver toxicity (ALT 4×ULN) receives BER −2 → normalized 0.00. The final score is (1.00 + 0.75 + 0.00)/3 = 0.58, demonstrating how safety signals directly penalize otherwise promising biomarker profiles.

This comprehensive methodology ensures that biomarker scoring reflects both therapeutic efficacy potential and safety risk assessment, weighted at 21% to significantly influence final asset prioritization while maintaining evidence-based rigor through mandatory citations.

#### 3.3.4 Development Reasons LLM

For all assets not currently FDA-approved, an automated investigator employs a multi-source intelligence gathering approach to determine development status and discontinuation rationale. **Step 1: Parallel Source Investigation** - The system simultaneously searches three primary information sources: company press releases, data collected from ClinicalTrials.gov registry for terminated/withdrawn/suspended trials with why stopped field analysis, and web news articles containing discontinuation announcements. Search queries target specific discontinuation keywords including “discontinued,” “terminated,” “FDA hold,” “clinical hold,” “suspended,” “halted,” while also detecting active development signals like “ongoing,” “enrolling,” **Step 2: LLM Analysis, Classification, and Evidence Validation** - GPT-4o analyzes aggregated findings from three parallel sources to classify the primary discontinuation reason into seven standardized categories with confidence scoring (0-100%): active development (1.0 score), active with safety concerns (0.7), business/operational reasons (0.6), unknown reasons (0.3), discontinued unclear why (0.1), efficacy failure (−0.2), and safety/regulatory discontinuation (−0.5). The LLM processes combined text from: (1) company press releases containing discontinuation announcements and regulatory actions, (2) ClinicalTrials.gov termination records with why stopped explanations, and (3) web news articles with fraud investigations, safety concerns, and development updates. Each classification includes supporting evidence with direct quotes, source citations, and detailed analysis summaries. The system specifically flags drugs under regulatory investigation, FDA holds, or with ongoing safety concerns even if still in development. **Step 3: Structured Output Generation** - Results are formatted into computable development status scores and reason classifications that directly feed into the downstream scoring algorithm, with 7-day caching to optimize API costs while maintaining current information.

#### 3.3.5 Ownership Intelligence

The ownership intelligence module employs a hybrid approach combining LLM-powered web search with structured database queries to build comprehensive asset ownership profiles. **Step 1: LLM Asset Owner Investigation** - GPT-4o with web search capabilities executes a systematic seven-point search strategy: ownership history queries, developer company identification, generic/brand name cross-referencing, licensing agreement searches, regulatory approval tracking across eight global markets (FDA, EMA, PMDA, NMPA, Health Canada, DCGI India, Roszdravnadzor Russia, MFDS South Korea), and specific indication mapping for each approved region. The LLM distinguishes between current asset owners (IP/development rights holders) and original developers, constructing complete ownership transfer chains. **Step 2: Clinical Trials Sponsor Analysis** - The system leverages sponsor metadata already collected during Stage 1 clinical trials data aggregation to identify recent trial sponsors and track sponsor changes over time for asset transfer detection. This analysis utilizes the existing sponsor history from the clinical trials database to extract lead sponsor information from active, recruiting, and completed studies, identifying the most recent sponsors and latest activity dates to validate the current ownership status without additional API queries. **Step 3: Patent Landscape Mapping** - When available, the system queries the PatentsView API (USPTO) and European Patent Office (EPO) Open Patent Services to identify primary patents, publication dates, applicant companies, and patent family relationships. This multi-jurisdictional analysis determines the freedom-to-operate status, patent cliff risks, and expiration timelines that affect commercial viability and licensing opportunities. All patent database queries are performed with appropriate rate limiting to ensure compliance with free API usage policies and to avoid service disruptions. **Step 4: Safety Signal Integration** - The drug safety client aggregates adverse event data from FDA FAERS, regulatory labels, and black-box warnings to complete the risk profile. This safety intelligence identifies withdrawn products, regulatory actions, and ongoing safety investigations that impact asset value and licensing feasibility.

The parallel execution of these five modules creates a comprehensive molecular, regulatory, and commercial characterization that forms the foundation for the subsequent scoring and ranking algorithms. All enrichment data is structured and stored in JSON format, with each candidate receiving a comprehensive profile containing normalized drug identifiers, ChEMBL/PubChem cross-references, target annotations, biomarker scores, regional approval mappings, development/discontinuation classifications, ownership intelligence, and patent landscapes. This standardized JSON schema enables efficient downstream processing, caching for performance optimization, and seamless integration with the scoring algorithms that consume these enriched candidate profiles.

### 3.4 Stage 3. Scoring and Prioritization

Following the enrichment stage, clinical-stage, not-yet-approved assets (both active and discontinued/shelved programs) undergo High Potential Scoring. FDA-approved drugs receive repurposing classification, and preclinical assets are propagated to final display.

The score, S, for each clinical-stage asset is a weighted sum of seven evidence-based components, normalized to a value between 0.0 and 1.0. The score is calculated using equation 2.

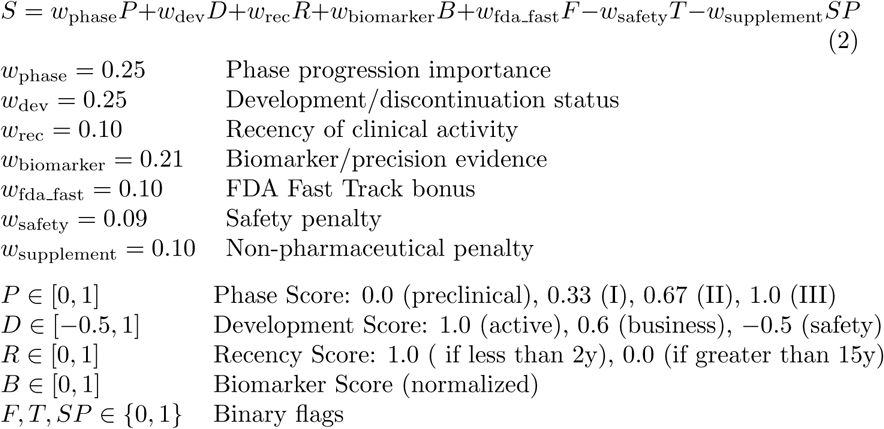

The weights in Equation 2 are expert-driven heuristics designed for a balanced, interpretable assessment. Their relative values are based on domain expertise, with clinical progression and development status weighted most heavily. The system’s transparency is a key feature, as shown in Figure 5, which allows for auditable,’white-box’ prioritization. Finally, the framework is designed to be flexible, as the weights can be adjusted to ‘reflect different prioritization strategies, allowing different stakeholders to tune the model to their specific needs.

**The Phase Score (P)** reflects the asset’s progression through clinical development, and its weight (w phase = 0.25) emphasizes the importance of successfully advancing through clinical trials. The scoring system awards 1.0 point for a drug if max phase is Phase III, 0.67 for Phase II assets, 0.33 for Phase I assets, and 0.0 points for Phase 0 or preclinical assets (no clinical progression). This linear progression scoring recognizes that each phase represents meaningful advancements in clinical validation, with later-stage assets receiving higher scores and a greater probability of regulatory success. **The Development Score (D)**, weighted at w dev = 0.25, directly measures the current development status of the asset as described in subsection 3.3.4. To incorporate quantitative evidence of drug performance, **the Biomarker Score (B)**employs a sophisticated, multi-step evidence-based assessment of drug performance on disease-relevant biomarkers, weighted at w biomarker = 0.21, as explained in subsection 3.3.3. **The Recency Score (R)**, weighted at wrec=0.10, serves as a proxy for continued investment by rewarding assets with recent clinical activity. A binary **FDA Fast Track Bonus (F)** (wfda fast =0.10) is awarded to assets that have gained regulatory recognition for their potential to address an unmet need. Finally, the model applies two penalties: a binary **Safety Penalty (T)**(wsafety = 0.09) is subtracted if any evidence of safety issues is detected, and a **Supplement Penalty (SP)** (wsupplement = 0.10) is applied to deprioritize supplements. These weights are heuristics-based and can be adjusted to reflect different prioritization strategies.

**Repurposing Opportunity Analysis:** Assets identified as FDA-approved but not indicated for the target disease are funneled into a separate analysis pipeline. For this cohort, an LLM evaluates each drug’s potential for repurposing by assessing its mechanistic fit for the target disease, reviewing any existing cross-indication clinical evidence, and considering its known safety profile. This process yields a list of repurposing candidates, each annotated with a supporting rationale.

### 3.5 Stage 4: Final Output and Provenance

The pipeline’s final output is a comprehensive JSON/pdf object containing the ranked lists of preclinical, high-potential discovery assets and repurposing opportunities. Each entry includes all enriched metadata, scores, discontinuation evidence, ownership status, and safety notes. Crucially, every data point is appended with provenance tags linking back to its original source document or API response, ensuring that all generated insights are transparent, traceable, and auditable for expert review.

## 4 Results

The LLM assisted pipeline was applied across three therapeutic domains: Alzheimer’s disease (AD), pancreatic cancer (PC), and cystic fibrosis (CF) to evaluate their drug landscape analysis. **Figure 1** provides an overview of the pipeline, which integrates multi-source data collection, enrichment, and categorization, followed by multi-criteria scoring and ranking of assets.

Across the case studies, the pipeline ingested a substantial evidence base of clinical trial data. **Figure 2** summarizes the landscape of registered clinical trials for each disease, using donut charts to represent trial status distributions. The detailed statuses from the registry (left panels) are consolidated into simplified ‘Main Categories’ (right panels) to facilitate high-level interpretation and downstream analysis. These main categories consist of ‘Completed’, ‘Active/Ongoing’, ‘Unknown/Other’, and ‘Failed/Stopped’, which aggregate statuses like ‘Terminated’, ‘Withdrawn’, and ‘Suspended’. Pancreatic Cancer had the largest volume of trials (N=6000) and the highest fraction of failed trials (13.1%) (See Figure2b). Alzheimer’s Disease (AD) had the second-largest volume of trials (N=2,449) and also a high proportion of failed trials (7.3%). By contrast, CF had a smaller absolute number of trials in the dataset and a relatively higher proportion of completed or active trials, reflecting recent successes in CF drug development (e.g., CFTR modulator therapies) and a more focused pipeline. The large “Failed” segments in AD and PC emphasize the high attrition rates in these diseases, reinforcing the importance of triaging resources toward candidates with the strongest supporting evidence. Developing a new drug typically spans ∼10–15 years and can require an investment of 1 billion dollars, underscoring the value of early identification of low-probability and high-potential projects. The breadth of trial records used here provides a robust foundation for scoring because it leverages longitudinal ClinicalTrials.gov evidence across each disease’s history [25, 26, 27, 28, 29].

**Figure 2:**
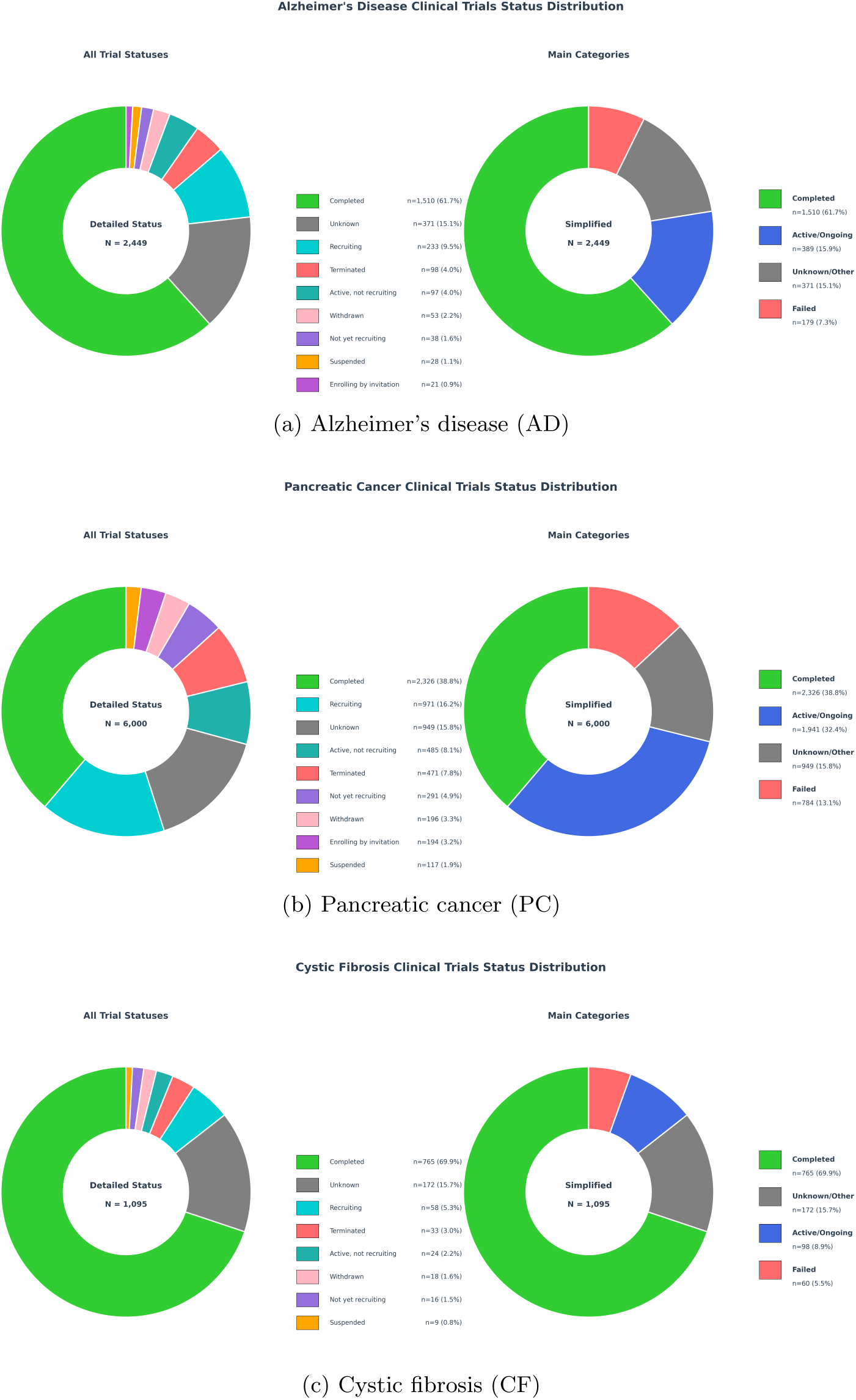
Clinical trial–status distributions across case-study diseases. (a) AD, (b) PC, (c) CF. ^15^

### 4.1 Scouting Score Reflects Evidence Accumulation

The initial triaging stage employs a Scouting Score, which integrates four key components: a Trial Points metric for trial volume, a Phase Bonus for developmental advancement, a phase-aware Failure Penalty, and a Total Reason Adjustment that differentiates between operational and scientific discontinuations (See subsection 3.2.1).

The Scouting Score effectively captures evidence accumulation, exhibiting a positive correlation with a drug’s total clinical trial activity across all three diseases, as shown in **Figure 3** (left panels; Pearson’s *r* = 0.611 for AD, 0.776 for PC, and 0.666 for CF). This relationship indicates that as a drug amasses more supportive data through clinical studies, its score increases. The scores also increase with each clinical phase, indicating support for late-stage programs (**Figure 3**, right panels). Additionally, Figure 3 demonstrates that an asset’s assigned clinical phase is not always proportional to its associated trial volume. For instance, several late-stage compounds, depicted as blue (Phase 4) data points in the left panels of Figure 3, are associated with a limited number of trials. This pattern frequently characterizes repurposed pharmaceuticals or supportive care agents that retain an advanced phase designation from a separate indication, despite being investigational for the disease in question. The presence of these assets also explains the significant score variance seen in the violin plots (**Figure 3**; right panel), as each phase category could contain a mix of both extensively-trialed and repurposed/supportive candidates.

**Figure 3:**
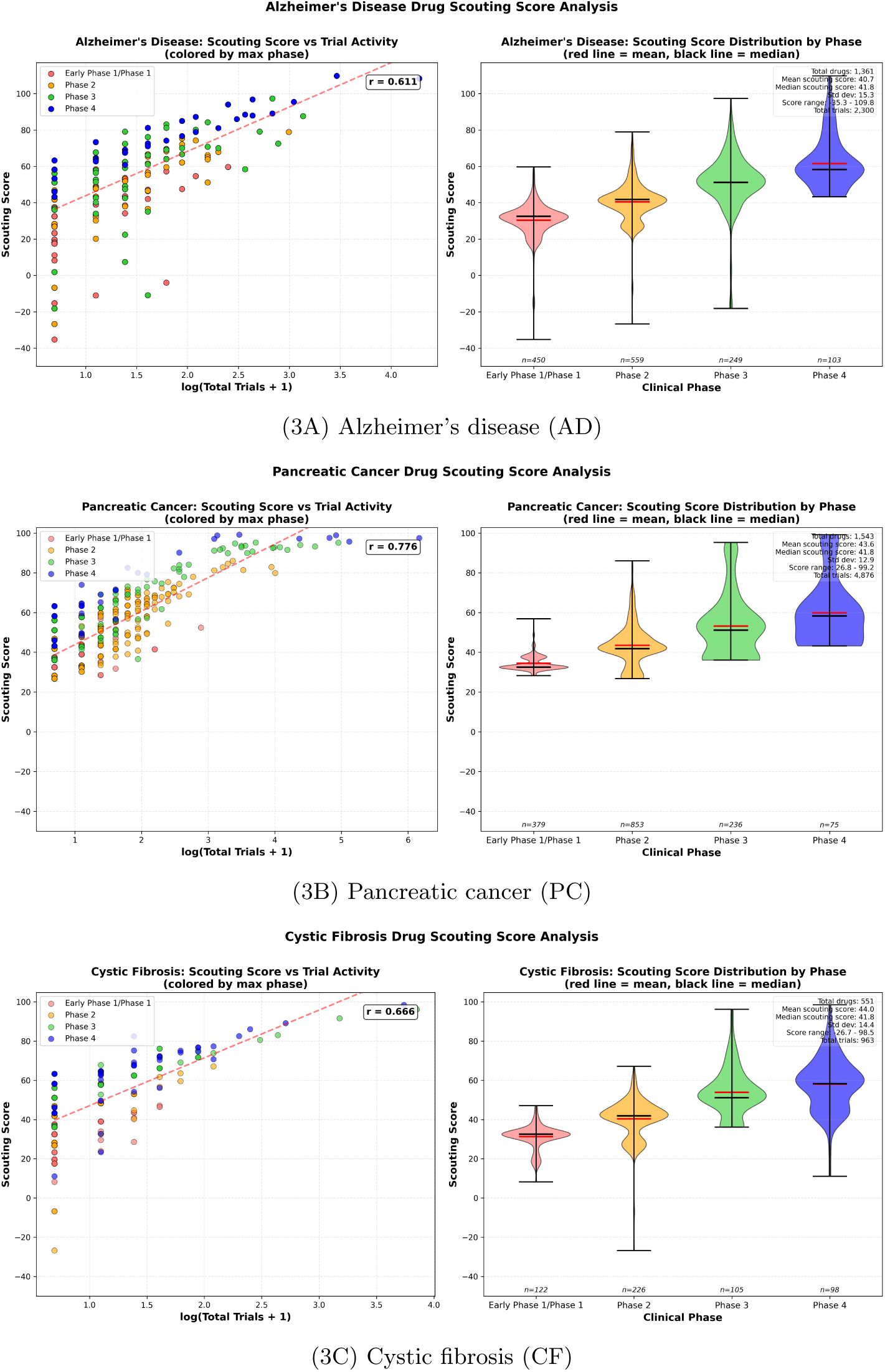
Scouting Score analysis across AD (3A), PC (3B), and CF (3C).

Figure 4 quantifies how the Total Reason Adjustment, Failure Penalty, and Phase Bonus components modulate the Scouting Score, with panels A–C corresponding to Alzheimer’s disease (AD), pancreatic cancer (PC), and cystic fibrosis (CF), respectively. The Total Reason Adjustment shows a positive association with the scouting score (Pearson’s *r* = 0.502 for AD, 0.438 for PC, and 0.219 for CF). This indicates that assets whose discontinuations are primarily for operational reasons (e.g., business or recruitment) rather than scientific ones (e.g., safety or efficacy) tend to earn higher scouting scores. In contrast, the Failure Penalty exhibits a negligible correlation in all three diseases (*r* = −0.013 for AD, −0.110 for PC, and −0.050 for CF), which is consistent with its role Scouting Score analysis across Alzheimer’s disease, pancreatic cancer, and cystic fibrosis. For each disease, the left panels show the relationship between total trial activity (log-scaled) and the computed Scouting Score, with points colored by highest clinical phase. Pearson’s r is shown for each correlation. The right panels display Scouting Score distributions across clinical phases (Unknown, I, II, III, IV); horizontal black lines indicate medians and red lines indicate means. Overall, Scouting Score increases with both trial volume and phase, reflecting progressive evidence accumulation, with tighter score variance at later stages as a phase-aware, Bayesian-smoothed tempering factor rather than a primary driver. The Beta(1,5) prior prevents the over-penalization of understudied drugs with limited trial data, ensuring that a single early failure does not disproportionately impact the scoring. Finally, the Phase Bonus correlates moderately with the score (r = 0.586 for AD, 0.568 for PC, and 0.677 for CF), reinforcing that later-stage programs are systematically scored higher. Whereas CF shows the strongest phase-score relationship (r = 0.677), likely reflecting the disease’s monogenic nature [30, 31, 32, 33] and the established CFTR modulation paradigm, where clinical phase advancement more reliably indicates therapeutic potential compared to the more complex pathophysiology and diverse mechanisms being pursued in AD and PC [34, 35]. Finally, the Phase Bonus positively correlates with the scouting score (*r* = 0.598 for AD, 0.568 for PC, and 0.677 for CF), reinforcing that later-stage programs are systematically scored higher.

**Figure 4:**
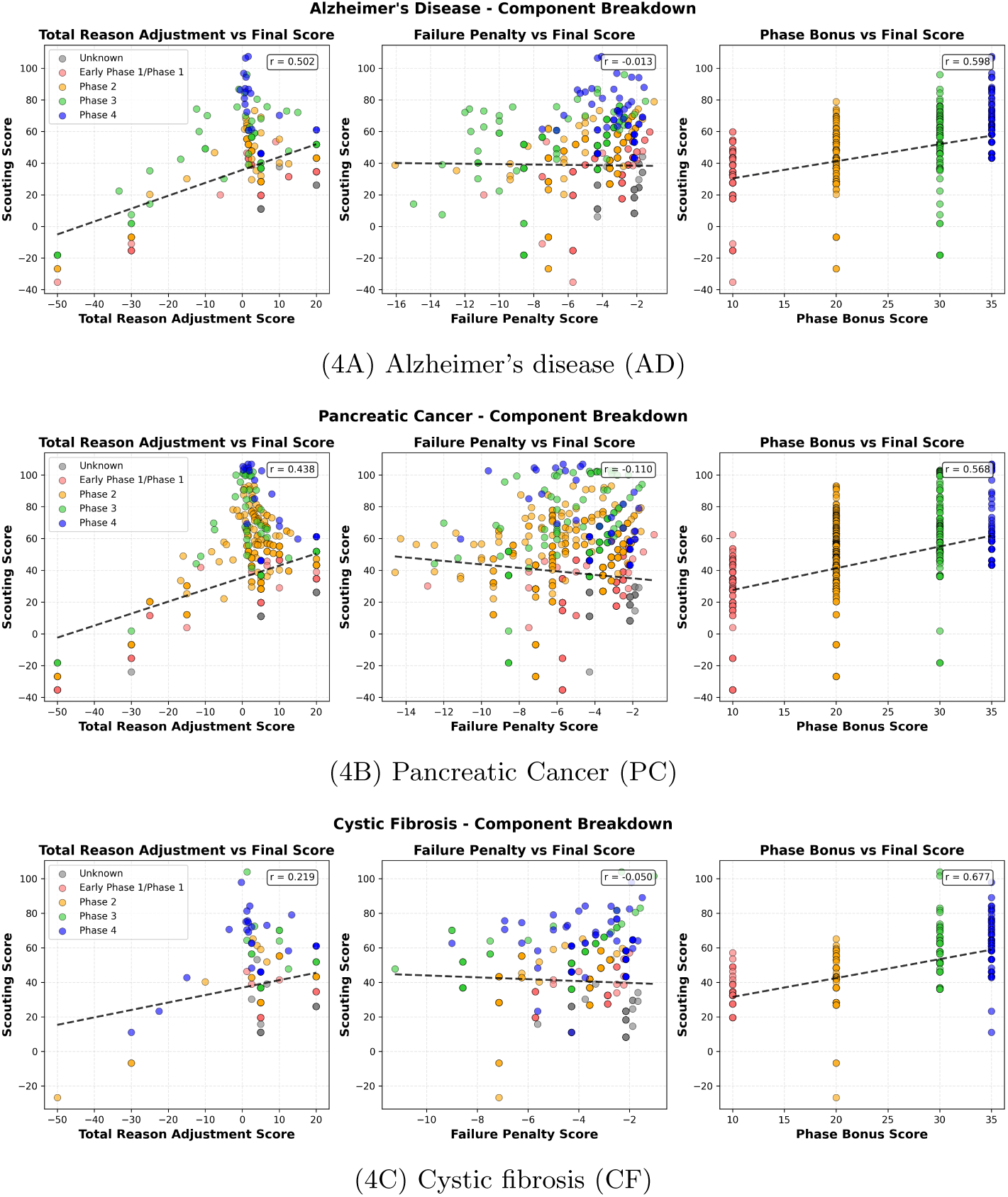
Component breakdown of the Scouting Score (*S*). Panels A–C correspond to Alzheimer’s disease (AD), pancreatic cancer (PC), and cystic fibrosis (CF). For each disease, scatter plots relate *S* to (left) Total Reason Adjustment, (middle) Failure Penalty, and (right) Phase Bonus; points are colored by maximum clinical phase and dashed lines indicate linear fits (insets report Pearson’s *r*). Total Reason Adjustment shows a positive association with *S*, whereas the Failure Penalty exhibits near-zero association by design—a phase-aware, Beta(1, 5)–smoothed tempering factor that avoids over-penalizing sparse programs. The Phase Bonus is moderately correlated with *S*, with CF showing the strongest phase–score coupling, consistent with its monogenic etiology and established CFTR-modulator paradigm.

Together, these results validate that the Scouting Score’s architecture behaves as intended. The strong correlation with trial volume (Figure 3) confirms that this metric acts as the score’s foundational driver, establishing a robust evidence baseline. The analysis in Figure 4 further demonstrates how factors such as phase advancement, a favorable discontinuation profile, and limited scientific failures act as key modulators that differentiate assets and fine-tune the final score.

### 4.2 Second-Stage Triaging: Multi-Modal Enrichment and Asset Categorization

After the Scouting Score calculation, the top 250 drugs are first classified by FDA approval status into three categories: approved for the same indication, approved for a different indication, or not approved. Subsequently, the drugs approved for a different indication and the not-approved drugs, along with pre-clinical and FDA-discontinued drugs, all enter the enrichment pipeline for comprehensive characterization.

In this enrichment process, each drug undergoes comprehensive characterization across multiple data modalities to build a complete evidence profile and enable refined prioritization, as described in the methods section 3.3. For example, Table 1 details the typical enrichment information for the top 4 Alzheimer’s drugs and the bottom 4 Alzheimer’s drugs.

**Table 1:**
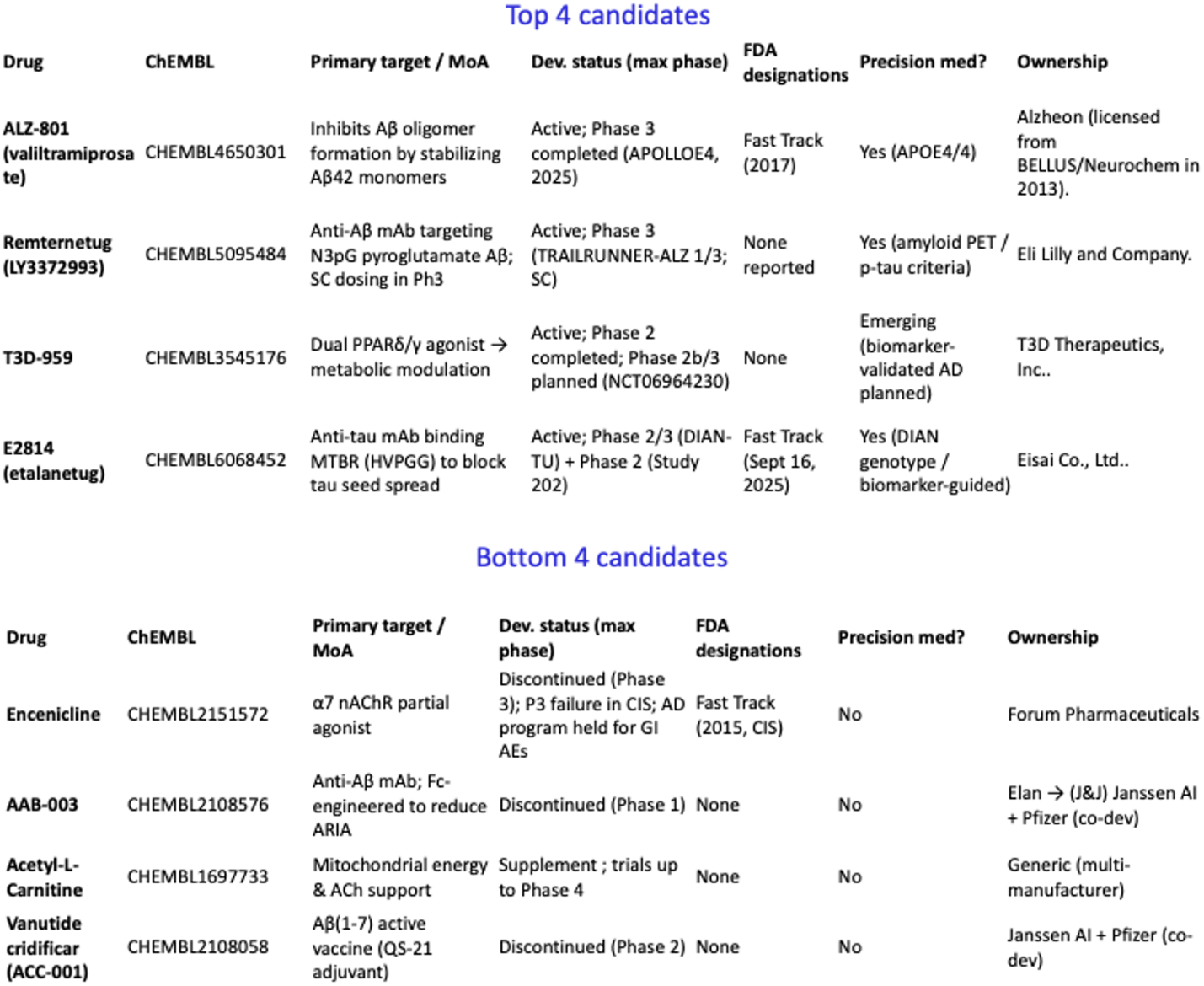
Comprehensive Multi-Modal Enrichment Profile for Top Investigational Alzheimer’s Disease Candidates.

The enrichment pipeline successfully characterized diverse therapeutic approaches spanning amyloid targeting, including oligomer inhibition with ALZ-801 [36, 37] and immunotherapy with Remternetug [38]. Additional mechanisms included metabolic modulation with the PPARδ/γ agonist T3D-959 [39], tau pathology via seed propagation blockade with E2814 [40], and cholinergic enhancement. Notable findings include the concentration of precision medicine approaches in top candidates and the FDA Fast Track designation for ALZ-801, indicating regulatory recognition of its therapeutic potential. While E2814 received FDA Fast Track designation in September 2025, all figures and data presented in this analysis were generated prior to August 2025, which explains why the FDA designation appears as “None” in our enrichment pipeline results.

Following enrichment, unapproved clinical-stage assets move into prioritization for weighted scoring and ranking (see Methods section 3.4), while repurposing candidates and preclinical assets are forwarded to the final output module.

Figure 5 shows the decomposition of score *S* into its weighted components for the top 15 and bottom 15 assets in each disease (A: AD, B: PC, C: CF). Across all three indications, top-ranked assets show large contributions from the Phase Score (green) and Development/Endpoint Status Score (blue), with frequent Biomarker contributions (purple) and occasional FDA Fast Track bonuses (yellow). In contrast, bottom-ranked assets are characterized by smaller Phase and Development components, sparse biomarker support, rare Fast Track bonuses, and, in several cases, visible negative penalties such as Safety/Regulatory (red) and/or Supplement (orange) that further depress totals. The pattern is consistent with the scoring design: the maturity of development and supportive clinical/biomarker evidence drive higher ranks, while scientific safety signals and non-pharmaceutical interventions are de-prioritized. In the Alzheimer’s disease (AD) chart Figure 5A, top-ranked assets are characterized by substantial Phase and Development contributions (green/blue bars), reflecting advanced clinical maturity. High scores are frequently augmented by a significant Biomarker component (purple bar), indicating linkage to translational endpoints [40],[41], [42]. Furthermore, several programs possess Expedited Program designations (yellow bar), such as the Fast Track status. This high-scoring profile is exemplified by assets including the Aβ-oligomer inhibitor ALZ-801 (valiltramiprosate) [37], the anti-Aβ N3pGlu mAb remternetug [38], the anti-tau MTBR mAb E2814/etalanetug [40], the PPAR-δ/γ agonist T3D-959 [39], the SIGMAR1 agonist blarcamesine (ANAVEX 2-73) [43], and the anti-Aβ mAb gantenerumab [44]. Additional programs with high scores, driven by late-phase data or robust translational packages, include GV-971 (sodium oligomannate) [45], CT1812 [46, 47], AR1001 [48], and BI 409306 [49]. Conversely, bottom-ranked AD assets (right panel, Figure 5A) display minimal Phase/Development components and sparse biomarker linkage, frequently coupled with penalties (red/orange bars). These penalties often correspond to programs terminated due to significant safety, tolerability, or regulatory challenges. Examples include the clinical hold for encenicline (gastrointestinal adverse events) [50], critical safety/tolerability issues (namely amyloid-related imaging abnormalities (ARIA-E) and microhemorrhage) observed for AAB-003 [51] and the γ-secretase inhibitor avagacestat, which was halted due to dose-limiting gastrointestinal and dermatologic toxicities, an increased risk of skin cancer, and a trend for cognitive worsening [52]. A lack of clinical efficacy is another primary contributor to low scores, as evidenced by failed trials for the Aβ vaccine ACC-001 (vanutide cridificar) [53], ponezumab [54], ELND005 (scylloinositol) [55], and dimebon (latrepirdine) [56]. Other low-scoring assets include those with mixed or negative randomized controlled Trial. (e.g., acetyl-L-carnitine [57]) or programs with otherwise limited or negative clinical data packages (such as AZD1446 [58], AZD5213 [59], and ispronicline [60]). Thus, the bar patterns for low-scoring assets confirm that limited clinical progression, failed efficacy readouts, or significant safety headwinds result in depressed (S) scores.

**Figure 5:**
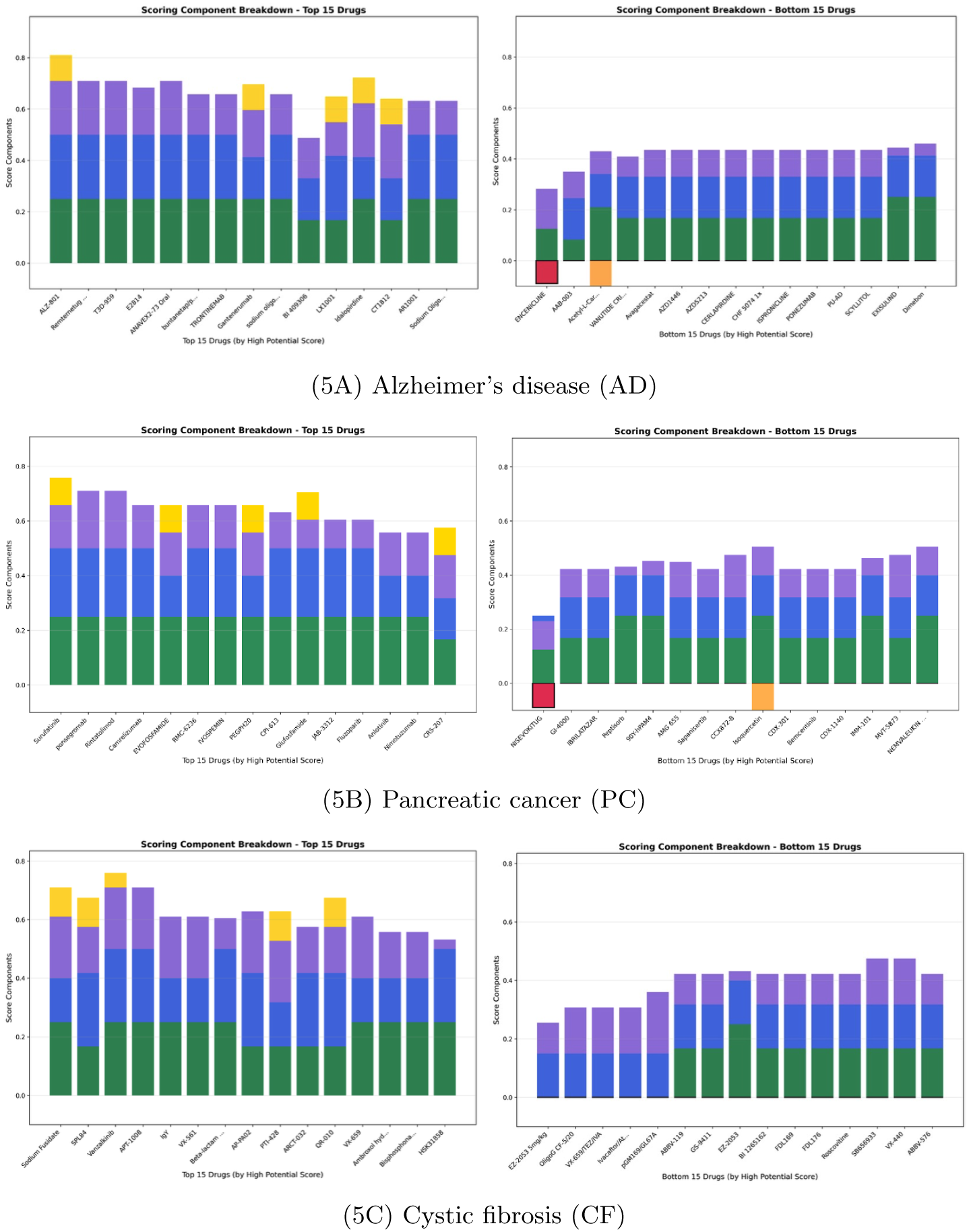
Component contributions to the Stage-3 score for the top and bottom 15 assets in each disease (A: AD, B: PC, C: CF). Top assets are dominated by Phase and Development/Endpoint contributions with added biomarker support and occasional FDA Fast Track bonuses; bottom assets show lower Phase/Development contributions and more frequent penalties (safety/regulatory, supplement).

For the Pancreatic Ductal Adenocarcinoma (PDAC) cohort (In the Alzheimer’s disease (AD) cohort (see Figure 5B), the Biomarker component (purple bar) is notably modest, even in top-ranked assets. This reflects the status of CA19-9, which is widely used for prognostic monitoring but is not a validated surrogate endpoint for survival. High scores are therefore primarily driven by Phase and Development/Endpoint data. Examples include surufatinib + camrelizumab combinations (NASCA) [61], the TLR3 agonist rintatolimod (Ampligen) in ongoing trials [62, 63], and the pan-RAS(ON) inhibitor RMC-6236 [64, 65]. This top-ranked group also includes several high-profile Phase 3 failures, such as the hypoxia-activated prodrug evofosfamide [66, 67], the stroma-targeting agent PEGPH20 [68], and the TCA inhibitor devimistat (CPI-613) [69], whose scores are driven by their advanced-stage data despite negative outcomes. Other assets, such as glufosfamide [70], JAB-3312 [71], anlotinib combinations [72], nimotuzumab [73], and the vaccine CRS-207 [74], further illustrate the microenvironmental barriers and biomarker sparsity that characterize the PDAC landscape.

Conversely, bottom-ranked PDAC assets exhibit minimal Phase/Development components and sparse biomarker linkage, with some incurring penalties (red/orange). This tier includes programs halted for futility or negative results, such as the KRAS vaccine GI-4000 [75] and the DR5 agonist conatumumab (AMG-655) [76]. It also includes investigational agents with mixed Phase 2 data (e.g., IMM-101 [77]), limited PDAC-specific activity (e.g., sapanisertib [78]), or early-phase immunotherapies (such as CDX-1140 [79] and CDX-301 [80]). Nutraceuticals like quercetin, which lack disease-modifying randomized controlled trials, also rank low and receive a **Supplement** penalty [81, 82]. Interestingly, the pipeline classified NIS793 (Nisevokitug, an anti–TGF-β antibody) under the Safety/Regulatory penalty after ingesting an external news report that an independent benefit–risk assessment in PDAC found higher mortality in the NIS793 arm versus the control [83]. Novartis then halted dosing/enrollment, discontinued development, and returned rights to XOMA, lowering the asset’s overall *S* score.

In the Cystic Fibrosis (CF) cohort (see Figure 5C), the prominent **Biomarker** signal (purple bar) in top-ranked assets reflects the field’s use of accepted CFTR response markers, such as sweat chloride and nasal potential difference (NPD), as established quantitative readouts of CFTR activity [84, 85, 86]. High-scoring examples exemplifying this pattern include the antisense oligonucleotide SPL84 [87], the CFTR amplifier nesolicaftor [88], the inhaled CFTR mRNA therapeutic ARCT-032 [89], and the ASO eluforsen (QR-010), which demonstrated human NPD and symptom improvements [90, 91].

Conversely, bottom-ranked assets feature smaller Phase/Development components and sparse biomarker linkage. This tier includes assets with limited or mixed efficacy signals, such as the alginate oligomer OligoG CF-5/20 [92] and the inhaled ENaC inhibitor BI 1265162, which proceeded through Phase 2 but failed to show a clinically relevant benefit [93]. Other programs were halted or superseded, including the CXCR2 antagonist SB-656933 (which showed biomarker shifts without clinical improvement) [94], the P2Y_2_ agonist denufosol (GS-9411) (Phase 3 negative on FEV_1_) [95], and the earlier-generation triple modulator regimen VX-659/TEZ/IVA [96]. Assets that entered clinical trials based on a strong preclinical rationale but ultimately showed no demonstrated clinical benefit, such as roscovitine (seliciclib) [97], also rank low. These bar patterns align with the scoring design: developmental maturity and robust CFTR-linked biomarker evidence elevate the (S) score, whereas a lack of late-phase efficacy or mixed readouts for non-modulator adjuncts assigns assets to the lower tier.

To validate the robustness of these assessments, we characterized the evidence base supporting each asset. Figure 6 summarizes the number of sources attached to each asset and the mix of source types for Alzheimer’s Disease (Fig. 6A), Pancreatic Cancer (Fig. 6B), and Cystic Fibrosis (Fig. 6C). Across indications, dossiers draw on roughly 9 – 14 sources per asset, with right-skewed tails for well-studied programs. Specifically, for Alzheimer’s Disease (Fig. 6A), there is a median of 12 and a mean of 12.4 sources per asset, with a source mix of 84.7 % scientific literature, 6.3 % curated databases, and 8.9 % industry news. For Pancreatic Cancer (Fig. 6B), the median is 11, and the mean is 14.1 sources per asset, comprising 87.5 % literature, 5.4 % databases, and 7.1% news. For Cystic Fibrosis (Fig. 6C), the median is 9, and the mean is 10.8 sources per asset, composed of 78.6 % literature, 8.6 % databases, and 12.7 % news. These distributions indicate that rankings are anchored in the literature and draw from multiple sources, with databases providing cross-checks and industry news supplying recent information.

**Figure 6:**
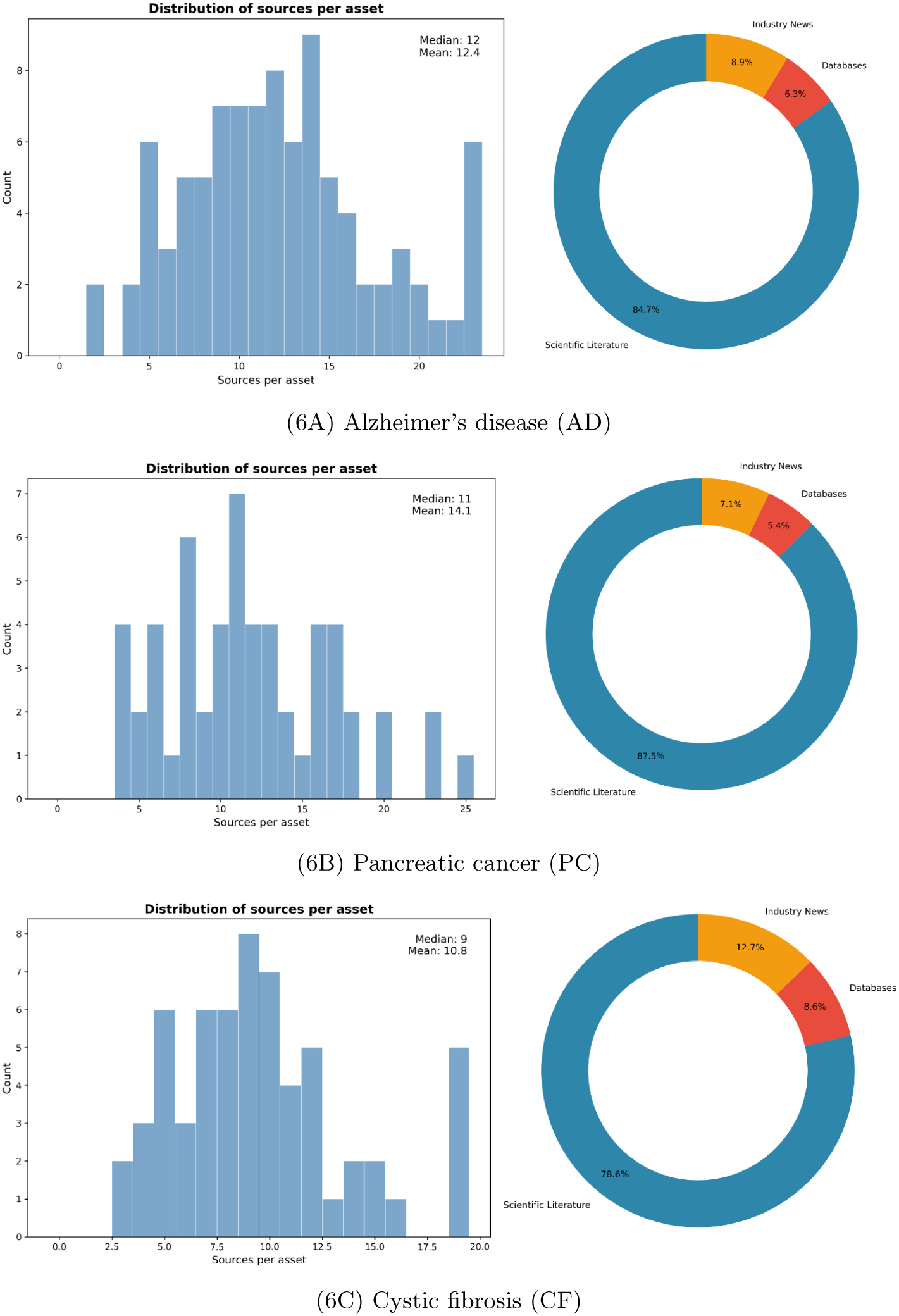
Evidence coverage per asset and source provenance. Histograms (left) show sources/asset (AD median 12/mean 12.4; PC 11/14.1; CF 9/10.8). Donuts (right) show source mix dominated by scientific literature (AD 84.7 %, PC 87.5 %, CF 78.6 %), with smaller contributions from databases and industry news

The final output of the triage process is a structured command-line interface (CLI) summary for each disease landscape. Figure 7, Figure S2, and Figure S3 show the command line interface displays for AD, PC, and CF, respectively. The output displays a Drug Classifications Summary, collapsing non-therapeutic items (e.g., diagnostic imaging agents and non-drug interventions) and flagged entries with significant safety/regulatory issues.

**Figure 7:**
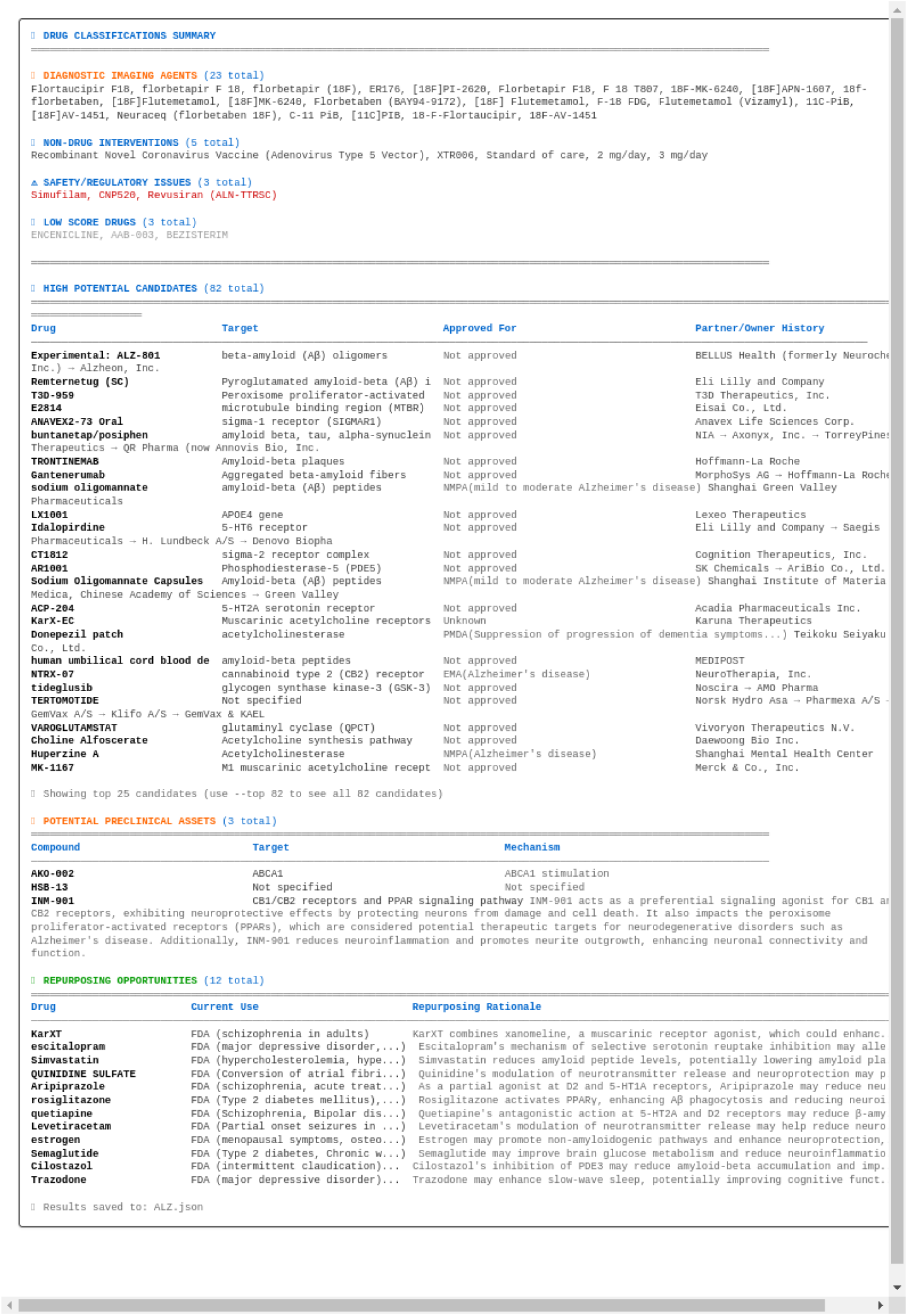
The command-line interface (CLI) summary for Alzheimer’s disease (AD) first filters non-therapeutic items and flags safety issues in its header. It then lists the top 25 (of 82) high-potential, not-yet-approved candidates, detailing their targets, global approval status, and owners. The summary concludes by enumerating 12 preclinical and repurposing candidates with rationales.

It then lists high-potential, not-yet-approved candidates (e.g., in the AD run, 82 total, with the top 25 displayed by default), with columns for drug, target/mechanism, and approval status across major regulators such as the EMA (EU/EEA), PMDA (Japan), NMPA (China), Health Canada (Canada), DCGI/CDSCO (India), Roszdravnadzor (Russia), and MFDS (South Korea), as well as ownership history.

Separate blocks enumerate potential preclinical assets, along with targets and mechanisms, as well as repurposing opportunities that are FDA-approved for other indications, each accompanied by a brief repurposing rationale. At the end of the run, the CLI saves the complete, provenance-linked dataset to JSON/pdf enabling auditability and downstream reuse. Data generated for Alzheimer’s is available in JSON format in the supplementary information.

The CLI includes a dedicated Safety/Regulatory Issues designation to flag assets with significant safety concerns or those under active regulatory investigation. This category includes programs such as simufilam, which is designated due to ongoing scientific scrutiny and public allegations documented in FDA citizen petitions and subsequent news coverage [98]. This designation is also applied to programs terminated for critical safety reasons. For example, CNP520 (umibecestat) was halted after interim analysis of the GENERATION prevention program revealed cognitive worsening in participants [99], a finding that was subsequently analyzed [100]. Similarly, the Phase 3 endeavor trial of revusiran was discontinued following the observation of a mortality imbalance [101, 102].

Although the pipeline is robust, several caveats motivate expert oversight to ensure the accuracy of regulatory and mechanistic data. A representative example appears in the Alzheimer’s disease output, where an asset labeled donepezil patch (see High Potential Candidates list in Figure 7) was incorrectly classified as a high-potential candidate, despite its FDA approval as ADLARITY on March 14, 2022 [103, 104, 105]. The donepezil transdermal system analysis illustrates a subtle but important limitation: when multiple pharmaceutical companies develop therapeutically equivalent formulations under different names, the system’s search methodology may create classification ambiguities that require human interpretation [106, 107].

Here, the pipeline correctly identified two distinct donepezil transdermal systems: (1) ADLARITY (Corium LLC; FDA-approved March 2022), and (2) Teikoku Seiyaku’s discontinued formulation (withdrawn from the U.S. market in 2012), as reflected in the JSON output reproduced below. However, the initial query for “donepezil patch” retrieved only the discontinued Teikoku product:

~~~
json
{
”drug name”: “Donepezil patch”,
”program status”: “development discontinued”,
”ownership chain”: “Teikoku Seiyaku Co., Ltd.”,
”discontinuation reason”: “development pause”,
”details”: “The donepezil patch has received approval in Japan and South Korea for the treatment of Alzheimer’s disease-related dementia. In the United States, the FDA issued a Complete Response Letter in 2011, and the application was subsequently withdrawn in 2012.”
}
While ADLARITY’s approval data was present in the regulatory database:
json { “drug”: “Donepezil”,
”fda approved”: true,
”filtered reason”: “fda approved drug”,
”ownership info”:
”regulatory”:
”orange book”:
”all products”: [
…, “trade name”: “ADLARITY”,
”trade name lower”: “adlarity”,
”ingredient”: “DONEPEZIL HYDROCHLORIDE”,
”dosage form route”: “SYSTEM;TRANSDERMAL”,
”approval date”: “Mar 11, 2022”,
”market status”: “RX”,
”applicant full name”: “CORIUM LLC”,
”nda number”: “212304”,
”rld”: “Yes”] }
~~~

This case demonstrates that generic therapeutic descriptors (”donepezil patch”) may not capture all relevant regulatory variants, particularly when approved formulations use proprietary brand names. Such scenarios underscore the necessity for human-in-the-loop validation to ensure comprehensive coverage of therapeutically similar compounds across different manufacturers, where name resolution can be confusing to Drug Resolver (see methods section), preventing potential oversights in the analysis in cases where multiple development pathways exist for identical therapeutic approaches.

A second caveat is exemplified by the preclinical asset HSB-13, which is displayed with its target and mechanism as “not specified” (Figure 7)). This annotation typically occurs when an asset’s development code is absent from curated repositories like ChEMBL, or when public disclosures are too limited or ambiguous for confident automated annotation. The case of HSB-13 highlights the value of a human-in-the-loop validation process. The pipeline’s LLM successfully identified a relevant source, as shown in the JSON snippet below (PMID: 20143421 [108]), but failed to extract actionable intelligence from it with sufficient confidence. This resulted in the “Not specified” annotations for its target and owner. However, a manual review of this exact same source revealed that HSB-13 is a multi-kinase inhibitor that targets GSK3, p38 MAPK, and cyclin-dependent kinases and has demonstrated neuroprotective efficacy in animal models [108]. The source also detailed its developmental history: HSB-13 originated from a collaboration between Southern Methodist University and UT Dallas before being licensed to EncephRx Inc., a startup focused on neurodegenerative therapeutics [109]. Public updates on HSB-13’s development have not been identified after 2011 (as of Oct 30, 2025) and its primary U.S. patent (US 8,680,094) is listed as “Expired – Fee Related,” lapsed for non-payment of maintenance fees [110](effective Mar 25, 2022). A manual search for HSB-13 also failed to yield a matching ChEMBL identifier corresponding to the name HSB-13.

While the system required manual intervention for a complex case like HSB-13, it is important to note its success in other instances. For example, it correctly identified other preclinical compounds, such as AKO-002 from Akome Biotech [111] and INM-901 by Inmed Pharmaceuticals [112], demonstrating its effectiveness as a primary triage and evidence-gathering tool.

~~~
{
   “drug_name”: “HSB-13”,
   “compound_name”: “HSB-13”,
   “development_stage”: “preclinical”,
   “source”: “llm_web_search_preclinical”,
   “verified_preclinical”: true,
   “clinical_trials_found”: 0,
   “raw_text”: “”,
   “licensing_available”: “unknown”,
   “indication”: “alzheimer’s disease”,
   “sponsor”: “Academic/Biotech”,
   “clinical_trials_count”: 0,
   “total_trials”: 0,
   “failed”: 0,
   “completed”: 0,
   “ongoing”: 0,    “phases”: [],
   “max_phase”: 0,
   “failure_ratio”: 0.0,
   “sponsors”: [   “Academic/Biotech”],
   “recent_active_trials”: 0,
   “sources”: “https://pubmed.ncbi.nlm.nih.gov/20143421/”
}
~~~

## 5 Conclusion

We present an AI-assisted triage pipeline that integrates clinical trials, regulatory records, and the scientific literature to build provenance-linked dossiers and an interpretable viability score for not-yet-approved drug assets, including active and shelved programs. Across Alzheimer’s disease, pancreatic cancer, and cystic fibrosis, the score increases with trial volume and clinical phase. Component analysis indicates that operational versus scientific discontinuations modulate but do not dominate rankings. The system is robust but requires human oversight. Regulatory name resolution can misclassify approvals (e.g., “donepezil patch” versus the FDA-approved donepezil transdermal system). Mechanism annotations may be missing when identifiers or disclosures are sparse (e.g., HSB-13). Curator-verified mappings, cross-regulator linkage, and the persistence of corrections should be applied. With these safeguards, the pipeline enables transparent and reproducible prioritization of clinical programs.

## Code Availability

All code and reproducible workflows are available at: github.com/anugrahat/asset*discoveryagent*.

## Notes

### Competing Interest Statement

The authors have declared no competing interest.

### Funding

- Startup Funds, Convexia, Felafax

