## Supplementary Figures for "Using Large Language Models to Assemble, Audit, and Prioritize the Therapeutic Landscape": Using-Large_Language_Models_to_Assemble__Audit__and_Prioritize_the_Therapeutic_Landscape__1_ 8-1.pdf

### Supplementary Information for: Using Large Language Models to Assemble, Audit, and Prioritize the Therapeutic Landscape

#### S1 Additional Methods

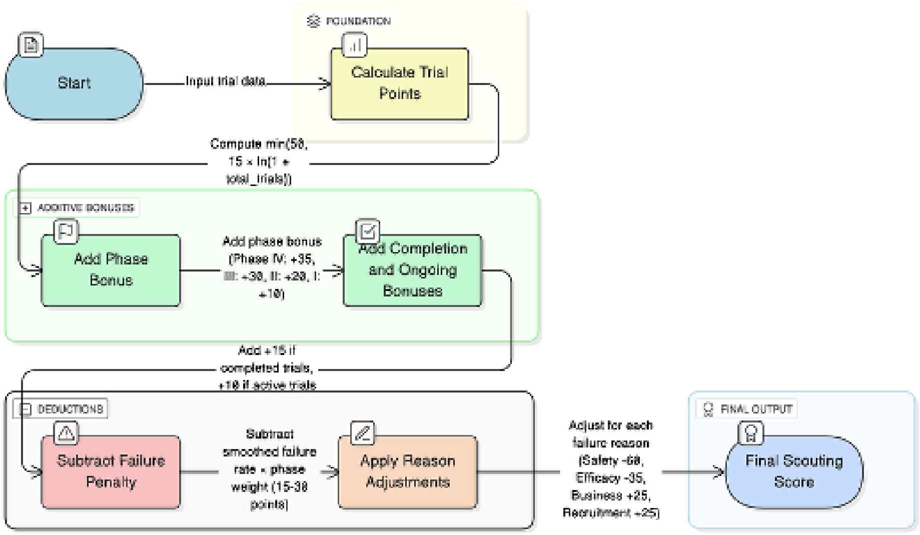


Figure S1: Scouting Score Calculation Workflow. The score combines a logarithmic Trial Points foundation (0-50) with bonuses for Phase (10-35), Completion (15), and Ongoing (10) trials. This is modified by a phase-weighted Failure Penalty and proportional Reason Adjustments (e.g., Safety: -60, Business: +25) based on LLM-classified termination reasons. The score ranks assets for enrichment (top 250).

```
DRUG CLASSIFICATIONS SUMMARY
════════════════════════════════════════════════════════════════════════════════

DIAGNOSTIC IMAGING AGENT (1 total)
18F-DOPA

NON-DRUG INTERVENTIONS (2 total)
GVAX Pancreas Vaccine, HyperAcute(R)-Pancreatic Cancer Vaccine

⚠ SAFETY/REGULATORY ISSUES (3 total)
Bardoxolone methyl, G17DT, dovitinib lactate

LOW SCORE DRUGS (2 total)
NISEVOKITUG, RP2D

════════════════════════════════════════════════════════════════════════════════

HIGH POTENTIAL CANDIDATES (52 total)
══════════════════════════════════════════════════════════════════════════════════════════════
════════════
Drug                          Target                              Approved For                            Partner/Owner History
──────────────────────────────────────────────────────────────────────────────────────────────────────────────────────────────────
Surufatinib                   VEGFR1, VEGFR2, VEGFR3, FGFR1, CSF  NMPA(Advanced extra-pancreatic neuroendocrine tumors...; Advanced
pancreatic neuroendocrine tumors (pNETs)) HUTCHMED has retained all rights to surufatinib worldwide since its development.
Folfirinox                    Thymidylate synthase, Topoisomeras  Unknown                                 Minneamrita Therapeutics
mFOLFIRINOX                   DNA synthesis and repair mechanism  Not approved                            Alliance for Clinical Trials in
Oncology
ponsegromab                   growth differentiation factor 15 (  Not approved                            Pfizer
Rintatolimod                  Toll-like receptor 3 (TLR-3)        Not approved                            AIM ImmunoTech (formerly
Hemispherx Biopharma)
Camrelizumab                  Programmed cell death protein 1 (P  NMPA(Hepatocellular carcinoma (second-line); Relapsed/refractory classic
Hodgkin's lymphoma ...; +6) Jiangsu Hengrui Pharmaceuticals Co., Ltd.
EVOFOSFAMIDE                  hypoxic tumor cells                 Not approved                            Threshold Pharmaceuticals → Merc
KGaA (co-development) → ImmunoGenesis
RMC-6236                      Oncogenic RAS mutations (G12X, G13  Not approved                            Revolution Medicines, Inc.
IVOSPEMIN                     polyamine biosynthesis enzymes (S-  Not approved                            Panbela Therapeutics, Inc.
IRGD                          αvβ3/5 integrins and neuropilin-1   Not approved                            Lisata Therapeutics, Inc.
High-dose                     Unknown                             Unknown                                 Fujian Shengdi Pharmaceu
Metabolic treatment           Unknown                             Not approved                            Health Clinics Limited
PEGPH20                       hyaluronan                          Not approved                            Halozyme Therapeutics, Inc.
CPI-613                       pyruvate dehydrogenase and α-ketog  Not approved                            Cornerstone Pharmaceuticals, Inc
→ Rafael Pharmaceuticals, Inc.
Glufosfamide                  DNA                                 Not approved                            Asta Medica (Degussa) → Baxter
International → Threshold Pharmaceuticals → Eleis
JAB-3312                      SHP2 (Src homology region 2 domain  Not approved                            Jacobio Pharmaceuticals
Fluzoparib                    PARP1 and PARP2 enzymes             NMPA(Ovarian cancer)                    Jiangsu Hengrui Medicine Co., Lt
Wild-type Reovirus            activated Ras signaling pathway     Not approved                            Oncolytics Biotech Inc.
Anlotinib                     Multiple tyrosine kinases includin  NMPA(locally advanced or metastatic NSCLC (NSCLC) after progression or
recurrence post at least two lines of systemic chemotherapy; soft tissue sarcoma; +4) CTTQ Pharma has maintained ownership since development
modified FOLFIRINOX regimen   DNA synthesis and repair mechanism  Unknown                                 Cornerstone Pharmaceutic
Nimotuzumab                   epidermal growth factor receptor (  NMPA(nasopharyngeal carcinoma; PDAC), DCGI(squamous cell carcinoma of the
head and neck) Tianjin Medical University Second Hospital
CRS-207                       mesothelin                          Not approved                            Aduro Biotech, Inc.
Huaier Granule                not specified                       NMPA(liver cancer)                      Qidong Gaitianli Pharmaceutical
Co., Ltd.
AMG 479                       insulin-like growth factor 1 recep  Unknown                                 NantCell, Inc.
CEA-targeted CAR-T cells      Carcinoembryonic antigen (CEA)      Unknown                                 Chongqing Precision Biot

Showing top 25 candidates (use --top 52 to see all 52 candidates)

POTENTIAL PRECLINICAL ASSETS (2 total)
════════════════════════════════════════════════════════════════════════════════
Compound                          Target                              Mechanism
────────────────────────────────────────────────────────────────────────────────
KAT 3-BP                          Hexokinase II (HK2)                 Inhibits glycolysis by covalently modifying hexokinase II,
leading to mitochondrial disruption and apoptosis in pancreatic cancer cells
THZ1                              Cyclin-dependent kinase 7 (CDK7)    THZ1 is a selective covalent inhibitor of CDK7, leading to
suppression of transcriptional activity and induction of apoptosis in pancreatic ductal adenocarcinoma (PDAC) cells. Its efficacy is
particularly notable in PDACs harboring the KRAS-G12V mutation, where it inhibits super-enhancer activity and the PI3K/AKT/mTOR signaling
pathway.

REPURPOSING OPPORTUNITIES (10 total)
══════════════════════════════════════════════════════════════════════════════════════════════
Drug                    Current Use                             Repurposing Rationale
──────────────────────────────────────────────────────────────────────────────────────────────
Oxaliplatin             FDA (adjuvant treatment of stag...)     Oxaliplatin forms DNA cross-links, inhibiting DNA replication and transcript
Capecitabine            FDA (Adjuvant treatment of pati...)     Capecitabine is metabolized to 5-fluorouracil, which disrupts DNA synthesis
Fluorouracil            FDA (Colorectal cancer, Esophag...)     Fluorouracil disrupts DNA synthesis by inhibiting thymidylate synthase, whic
Cisplatin               FDA (testicular cancer, ovarian...)     Cisplatin forms covalent bonds with DNA, leading to crosslinks that interfer
Pembrolizumab           Unknown                                As an immune checkpoint inhibitor, Pembrolizumab enhances T-cell mediated im.
Bevacizumab             FDA (Metastatic colorectal canc...)     Bevacizumab inhibits angiogenesis by blocking VEGF, which is crucial for tum
Nivolumab               FDA (Unresectable or metastatic...)     Nivolumab, as an immune checkpoint inhibitor, could enhance anti-tumor immun
Cetuximab               Unknown                                Cetuximab targets EGFR, which is often overexpressed in pancreatic cancer, p.
Docetaxel               FDA (locally advanced or metast...)     Docetaxel stabilizes microtubules and induces apoptosis, which could be effe
Carboplatin             FDA (ovarian cancer, lung cance...)     Carboplatin forms DNA cross-links, disrupting replication and transcription,

Results saved to: pan.json
```

Figure S2: Supplementary CLI summary for pancreatic cancer (PDAC): drug classifications and safety/regulatory flags; high-potential not-yet-approved candidates with target/MoA, global approval status, and ownership; plus blocks for potential preclinical assets and repurposing opportunities.

```
DRUG CLASSIFICATIONS SUMMARY

NON-DRUG INTERVENTIONS (1 total)
Standard of care IV antibiotic(s)

⚠ SAFETY/REGULATORY ISSUES (1 total)
Evening Primrose Oil

LOW SCORE DRUGS (5 total)
VX-659/TEZ/IVA, Ivacaftor/Ataluren, pGM169/GL67A, EZ-2053, OligoG CF-5/20

HIGH POTENTIAL CANDIDATES (56 total)

Drug                        Target                        Approved For                          Partner/Owner History

Sodium Fusidate             Staphylococcus aureus         EMA(skin infections caused by susceptible bacteria), PMDA(skin infection
caused by susceptible bacteria), NMPA(skin infections caused by susceptible bacteria), HC(skin infections caused by susceptible bacteria),
DCGI(skin infections caused by susceptible bacteria) Aceragen
SPL84                       CFTR mRNA with 3849+10 kb C->T mut  Not approved                    SpliSense
Vanzalkinib                 Cystic fibrosis transmembrane cond  Unknown                         Unknown
APT-1008                    dietary fats, proteins, and carboh  Unknown                         Forest Laboratories
IgY                         Pseudomonas aeruginosa bacteria     Unknown                         Blokhin's Russian Cancer
VX-561                      CFTR protein                  Not approved                          Concert Pharmaceuticals → Verte
Pharmaceuticals
Beta-lactam antibiotic      bacterial cell wall synthesis Unknown                               Fondation Hôpital Saint-
AP-PA02                     Pseudomonas aeruginosa        Not approved                          Armata Pharmaceuticals
PTI-428                     Cystic Fibrosis Transmembrane Cond  Not approved                    Proteostasis Therapeutics, Inc.
ARCT-032                    Cystic Fibrosis Transmembrane Cond  Not approved                    Arcturus Therapeutics
QR-010                      CFTR mRNA with F508del mutation     Not approved                    ProQR Therapeutics N.V.
VX-659                      CFTR protein                  Not approved                          Vertex Pharmaceuticals
Ambroxol hydrochloride 30 mg  Cystic fibrosis transmembrane cond  EMA(acute and chronic bronchopulmonary diseases ass...), PMDA(acute and
chronic bronchopulmonary diseases ass...), NMPA(acute and chronic bronchopulmonary diseases ass...), DCGI(acute and chronic bronchopulmonar
diseases ass...) Zambon SpA
Bisphosphonate treatment    Osteoclasts                   Unknown                               Merck Sharp & Dohme LLC
HSK31858                    unknown                       Not approved                          Haisco Pharmaceutical Group Co.,
Ltd.
CTX-4430                    Leukotriene A4 Hydrolase (LTA4H)    Not approved                    Estrellita Pharmaceuticals, Inc
Celtaxsys, Inc.
VX-522                      Cystic Fibrosis Transmembrane Cond  Not approved                    Vertex Pharmaceuticals
Liprotamase                 digestive enzymes lipase, protease  Not approved                    Anthera Pharmaceuticals
CSL787                      unknown                       Not approved                          CSL Behring
Ataluren                    CFTR protein                  Not approved                          PTC Therapeutics
Andecaliximab               Matrix metalloproteinase-9 (MMP-9)  Not approved                    Gilead Sciences, Inc. → åshibio
Liprotamase Powder for Oral S  Dietary fats, proteins, and carboh  Not approved                 Anthera Pharmaceuticals
PBI4050                     GPR40 agonist and GPR84 antagonist  Not approved                    ProMetic Life Sciences Inc.
alginate oligosaccharide    Mucus and bacterial biofilms in th  Not approved                    AlgiPharma AS
PTI-801                     CFTR protein                  Not approved                          Proteostasis Therapeutics, Inc.

Showing top 25 candidates (use --top 56 to see all 56 candidates)

POTENTIAL PRECLINICAL ASSETS (2 total)

Compound                    Target                        Mechanism

NBD1                        stabilizers of NBD1 protein   Unknown
NEU1 Inhibitor              Neuraminidase-1 (NEU1)        Inhibits NEU1 to reduce desialylation and shedding of mucin-1
ectodomain, thereby improving mucociliary clearance and reducing pulmonary inflammation and fibrosis

REPURPOSING OPPORTUNITIES (7 total)

Drug             Current Use                            Repurposing Rationale

Azithromycin     FDA (acute bacterial exacerbati...)    Azithromycin may induce overexpression of MRP, enhancing chloride conductanc
Rifampin         FDA (tuberculosis, leprosy, Hae...)    Rifampin inhibits bacterial RNA synthesis, which could potentially enhance t
Sildenafil       FDA (erectile dysfunction, pulm...)    Sildenafil, as a PDE5 inhibitor, may improve pulmonary function in cystic fi
Doxycycline      FDA (Rocky Mountain spotted fev...)    Doxycycline inhibits MMPs, which may reduce airway inflammation and tissue r
Roflumilast      FDA (chronic obstructive pulmon...)    Roflumilast, as a PDE4 inhibitor, may reduce inflammation in the lungs, whic
Insulin          FDA (Type 1 diabetes, Type 2 di...)    Insulin therapy is essential for managing cystic fibrosis-related diabetes (
Colistin         FDA (infections caused by multi...)    Colistin is effective against multidrug-resistant bacteria, which are common

Results saved to: cf.json
```

Figure S3: Supplementary CLI summary for cystic fibrosis (CF): drug classifications and safety/regulatory flags; high-potential not-yet-approved candidates with target/MoA, global approval status, and ownership; plus blocks for potential preclinical assets and repurposing opportunities.
